# Quartile coefficient of variation is more robust than CV for traits calculated as a ratio^1^

**DOI:** 10.1101/2022.10.13.512014

**Authors:** Zoltán Botta-Dukát

## Abstract

Comparing within-species variations of traits can be used in testing ecological theories. In these comparisons, it is useful to remove the effect of the difference in mean trait values, therefore measures of relative variation, most often the coefficient of variation (CV), are used. The studied traits are often calculated as the ratio of the size or mass of two organs: e.g. SLA is the ratio of leaf size and leaf mass. Often the inverse of these ratios is also meaningful; for example, the inverse of SLA is often referred to as LMA (leaf mass per area). Relative variation of a trait and its inverse should not considerably differ. However, it is illustrated that using the coefficient of variation may result in differences that could influence the interpretation, especially if there are outlier trait values. The alternative way for estimating CV from the standard deviation of log-transformed data assuming log-normal distribution and Kirkwood’s geometric coefficient of variation free from this problem, but they proved to be sensitive to outlier values. Quartile coefficient of variation performed best in the tests: it gives the same value for a trait and its inverse and it is not sensitive to outliers.

## 1. Introduction

Values of qualitative traits can considerably vary among and within species ^1,2^. The structure of variation can be explored by partitioning total variance into components related to different sources (e.g. variation between species, between sites within species, between individuals within site, and within individuals). The calculated variance components express the relative contribution of sources in percentage, therefore their values can be compared between sites, species, or traits. However, some ecological hypotheses are related to the extent of trait variation ^1,2^. For example, it is hypothesized that the extent of intraspecific trait variation is higher in generalists than in specialist species ^3,4^, and it may change along environmental and species richness gradients ^5–7^.

The coefficient of variation (CV), the standard deviation divided by the arithmetic mean, is the most widely used measure of the extent of trait variation ^e.g. 8–11^. CV has two advantages: it is a dimensionless measure of relative variation ^2^. The extent of trait variation can be compared among traits only if it is measured in the same units. For example, the standard deviation of height, measured in cm, and SLA, measured in g cm^−2^ cannot be compared, while their CV is comparable because it is dimensionless. Comparing absolute variation of the same trait between species also can be misleading when the difference between means is large. Ten centimeters departure from the mean height of the species is large for a short forb but small for a tall tree. That is why better to use relative measures, such as coefficient of variation, for among-species comparisons too.

Several papers called the attention to cases where CV should not be used ^e.g. 12,13^. The most important restrictions are that the domain of the variable has to be non-negative (otherwise, its arithmetic mean could be zero preventing calculation CV) and it has to be measured in ratio or log-interval scale, where the meaning of “zero” value is unarbitrary. It cannot be calculated for nominal or ordinal scale data, where the mean and standard deviation is undefined. CV also should not be calculated for interval and difference scale ^i.e. for log-transformed ratio-scale variables;,14^, where changing of unit influences the mean value. Brendel ^15^ pointed out that the CV of standardized stable isotope ratios depends on the applied reference isotope ratio. The aims of this paper are (1) calling attention to another problem: swapping nominator and denominator of ratio type traits results in an altered CV value; and (2) suggesting to use of quantile coefficient of variation that is free from this problem.

Ratios of size or mass of plant organs are widely used as functional traits, such as the ratio of leaf area and leaf dry mass (specific leaf area, SLA or leaf mass per area, LMA), the ratio of root length and root dry mass (specific root length, SRL) or ratio of the shoot and root mass ^16^. In these ratios, nominator and denominator are often interchangeable without loss of meaning; for example, instead of the specific leaf area (SLA) often its inverse, the leaf mass ratio (LMA) is calculated ^17^. We would expect the relative variation of the two forms of ratio (e.g. SLA and LMA) to be the same.

## 2. Theory

The coefficient of variation is defined as the ratio of standard deviation and mean of the distribution:

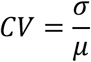

Regarding the ratio of two random variates to bivariate function allows approximating its mean and standard deviation by Taylor series expansion (see Appendix A for the derivation of formulas):

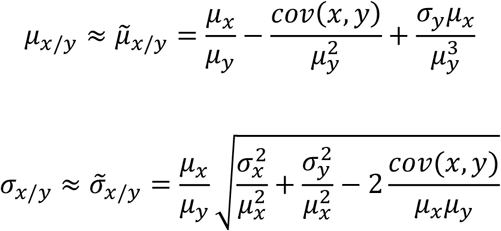

If CVs of *x*/*y* and *y*/*x* equal:

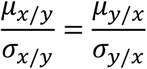

Therefore

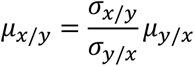

This equation should be - at least approximately – hold to approximate means and standard deviations, but:

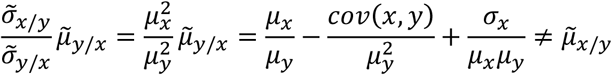

Since the equation does not hold for the approximate value, we can expect that CVs of a ratio and its inverse may differ. A real example will be shown in the Results section to illustrate that the difference could be important.

However, there is an important exception, when the ratio follows log-normal distribution. If x/y is log-normally distributed, its logarithm follows normal distribution, with *ν* mean and *θ* standard deviation

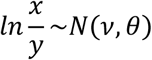

where *ν* and *θ* are the mean and standard deviation of the log-transformed ratio, respectively. The mean and standard deviation of the log-normal distribution are

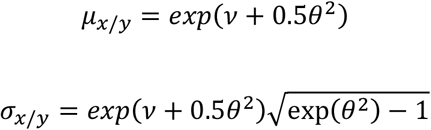

Therefore, CV depends on *θ* only, and it is independent from *ν* ^18^:

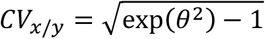

Since

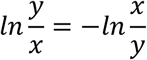

the logarithm of the inverse ratio is also normally distributed with the same standard deviation:

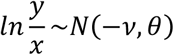

thus in this case CV is the same for the ratio and its inverse.

CV can be estimated by replacing standard deviation (*σ*) and means (*µ*) with their estimates (s and m, respectively):

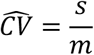

If x/y follows lognormal distribution, there is another estimator of CV:

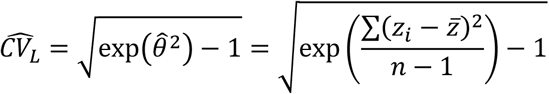

where 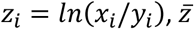 is the arithmetic means of log-transformed ratios and *n* is the sample size. 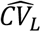 can be used as a descriptive statistic even if the ratio does not follow log-normal distribution. Kirkwood ^19^proposed another descriptive statistic the so-called geometric coefficient of variation:

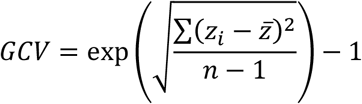

GCV is not an estimate of CV, even if *z* follows log-normal distribution.

The logic of calculating CV is that dividing the measure of dispersion (standard deviation in CV) by the measure of location (mean in CV) removes the effect of differences in dispersion due to different locations, and if both are measured in the same units results in a dimensionless measure. Following this logic, several alternatives to CV were developed. The main motivation was to develop more robust (i.e. less sensitive to outlier values) alternatives to CV^20 and references therein^. Unfortunately, most of the proposed robust relative variation measures are also sensitive to swapping nominator and denominator in ratio type traits. An exception is the quartile coefficient of variation (CV_Q_):

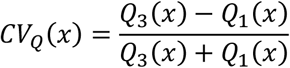

where *Q*_1_(*x)* and *Q*_3_(*x)* are the first and third quartiles of variable *x* ^20,21^.

For proving that *CV*_*Q*_(*x*/*y)* = *CV*_*Q*_(*y*⁄*x)* we will use the equation *Q*_3_(*y*/*x)* = 1⁄*Q*_1_(*x*/*y)*. Therefore, first, this equation has to be proved. Let us start from the definition of first

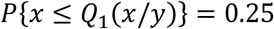

and third quartile

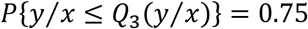

From the definition of the first quartile of x

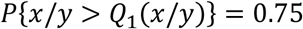

thus

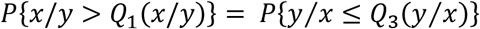

If *x*/*y* > *Q*_1_(*x*/*y)* then *y*⁄*x* < 1⁄*Q*_1_(*x*/*y)*, therefore

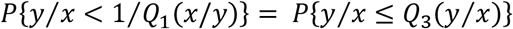

Since for a continuous variable, the probability of any possible value is zero, on the right side the “less than or equal to” can be replaced by “less than”

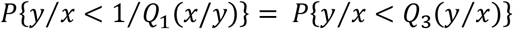

and this equation holds only if

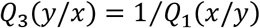

Now, we can turn back to the proof of *CV*_*Q*_(*x*/*y)* = *CV*_*Q*_(*y*⁄*x)* equality.

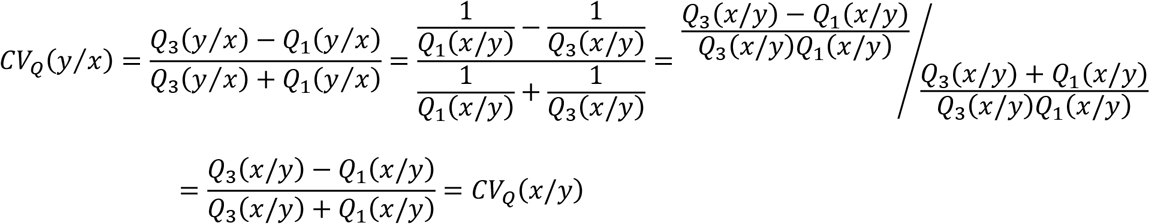

Note that finite sample estimates of *CV*_*Q*_(*y*⁄*x)* and *CV*_*Q*_(*x*⁄*y)* may slightly differ.

## 3. Results

As expected, the differences between SLA and LMA in 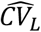 and GCV came only from rounding errors: the order of largest difference was 10^−16^. In the quartile coefficient of variation, the highest difference was 0.007 (Figure 1.a). However, differences hardly influenced the ranking of species according to the amount of intraspecific trait variation: the largest difference in ranks was 1, and 67 of 79 species the rank was the same for both traits. However, the amount of ITV of SLA and LMA measured by 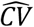 (i.e. estimated standard deviation divided by sample mean) differed considerably (Figure 1.b): the largest difference was 1.07. Although the rank of species based on SLA and LMA was strongly correlated even if ITV was measured by 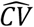 (Figure 2), the position of some species was strongly influenced: the largest difference in ranks between the two traits (SLA and LMA) was 21, and only 4 of 79 species remained ranks the same.

**Figure 1:**
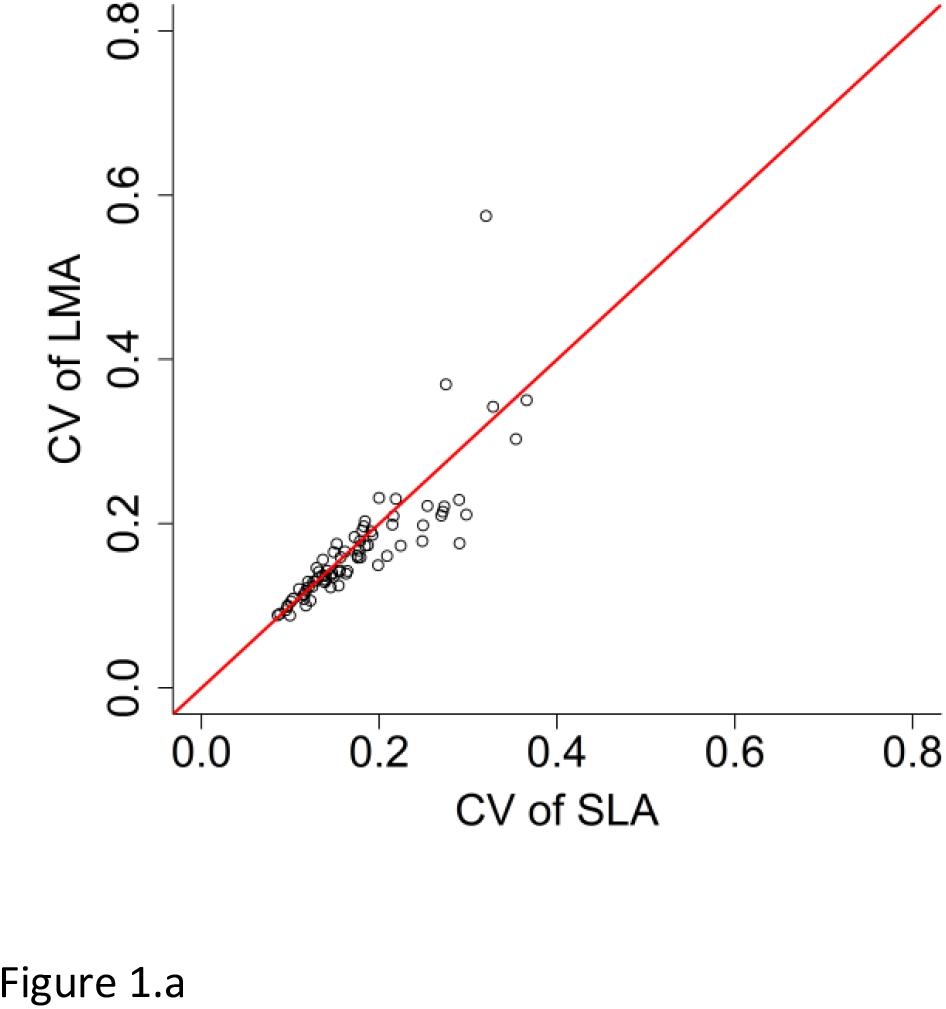
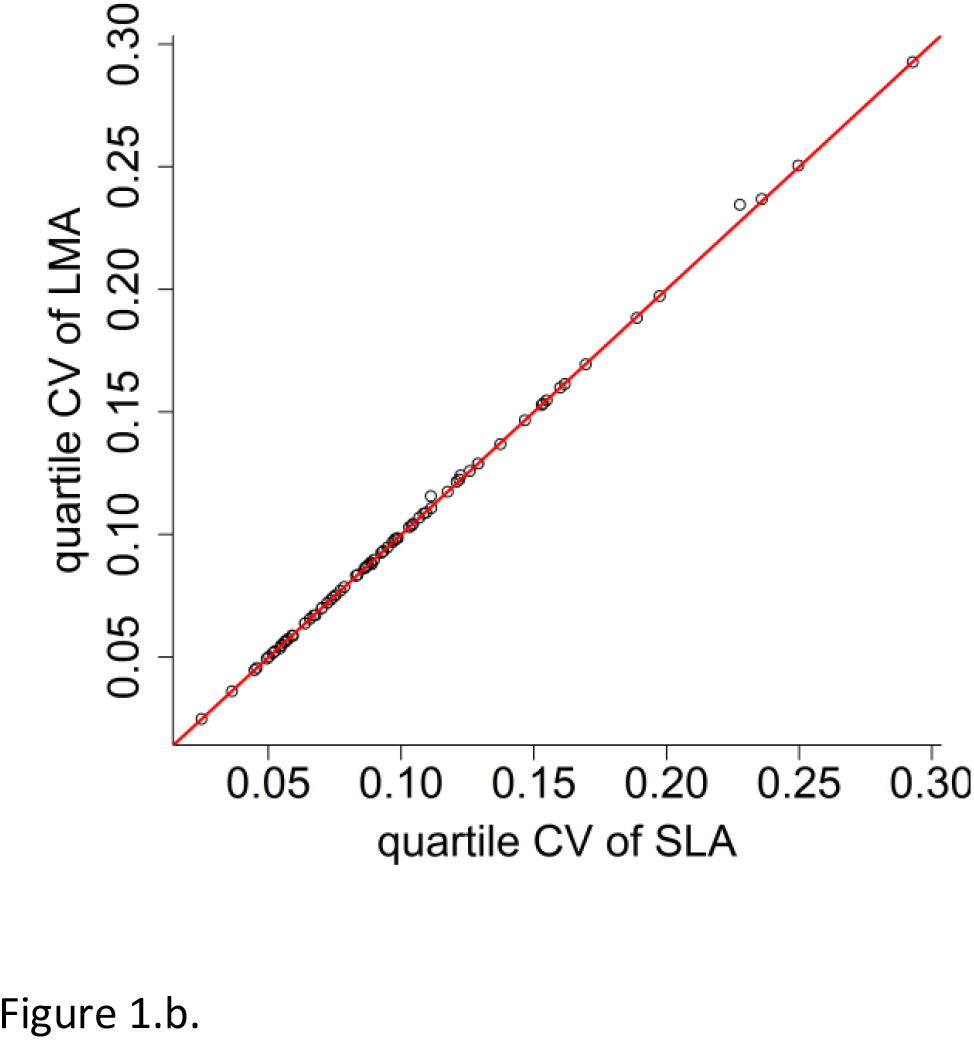
Within-species relative variation of specific leaf area (SLA) and leaf mass per area (LMA) calculated by (a) CV (coefficient of variation, standard deviation divided by mean) and (b) quartile coefficient of variation (see formula in the main text). Red line is the 1:1 line.

**Figure 2:**
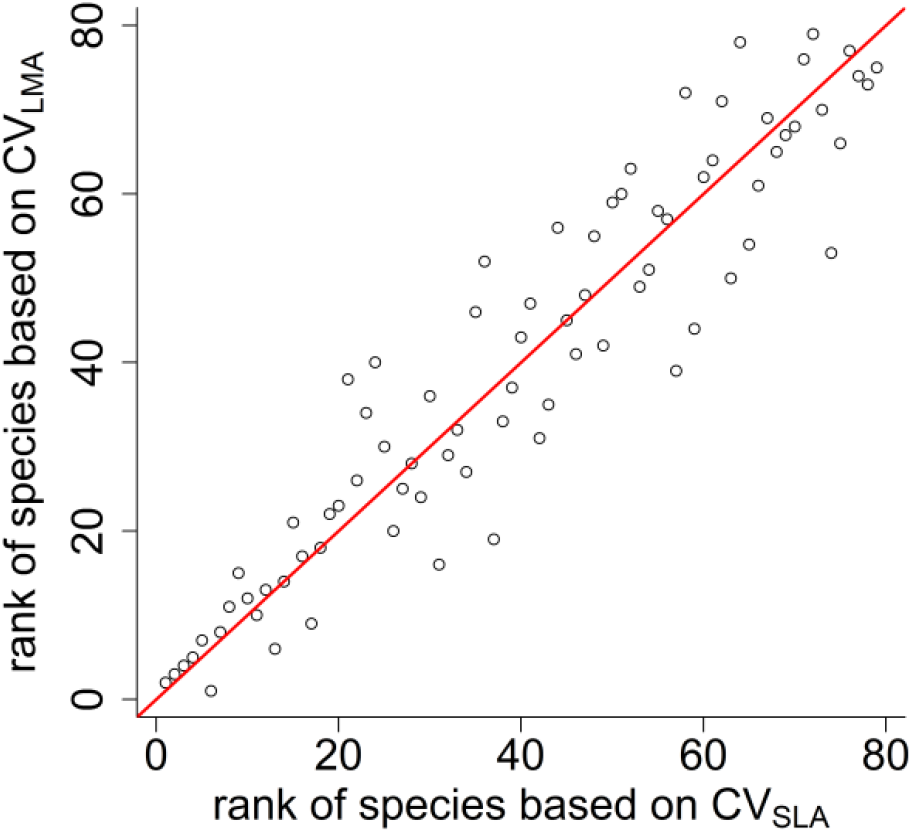
Rank of species based on their within-species relative variation of specific leaf area (SLA) and leaf mass per area (LMA) calculated by CV (coefficient of variation, standard deviation divided by mean). Red line is the 1:1 line.

The differences in 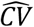 between SLA and LMA were mainly caused by outlier values. After species-wise excluding outlier SLA values, the highest difference reduced to 0.25, but the difference between ranks of species according to ITV of SLA and LMA remained large: the highest rank difference was 24 (even larger than without excluding outliers), and only for 14 of 79 species were the two ranks the same.

Excluding outlier values had a negligible effect on ITV measured by quadratic CV, the correlation between values estimated with and without excluding outliers was 0.99. The same correlation of 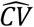 was 0.84. Surprisingly, the correlations between ITV calculated with or without excluding species-wise outliers were even smaller for 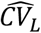 and *GCV* (0.67 and 0.65, respectively).

All of the four measures of ITV indicate almost the same property of species (Table 1): the lowest linear correlation was 0.61, while the lowest Spearman’s rank correlation was 0.72. Quartile coefficient of variation was the most different from the other three measures because it depends only on the central part of trait distribution, and therefore it is fully insensitive to outlier values.

**Table 1:** Correlations between within-species relative variation of SLA with (upper half-matrix) and without (lower half-matrix) excluding outliers.

| | $\widehat{CV}$ | $\widehat{CV}_L$ | $GCV$ | $CV_Q$ |
| --- | --- | --- | --- | --- |
| $\widehat{CV}$ | | 0.982 | 0.982 | 0.679 |
| $\widehat{CV}_L$ | 0.819 | | 1.000 | 0.718 |
| $GCV$ | 0.797 | 0.999 | | 0.718 |
| $CV_Q$ | 0.765 | 0.629 | 0.609 | |

## 4. Discussion

Presented results illustrate that ratio of sample standard deviation and sample mean 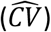 is sensitive both to outlier values and choosing a ratio-trait or its inverse (for example SLA or LMA). Three alternatives to this measure were evaluated in this paper. Both 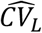 and GCV gave the same value for a trait and its inverse, but they are more sensitive to outlier values than 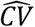. Quadratic CV proved to be the most robust measure of ITV, it was hardly influenced by either excluding outliers and choosing a trait or its inverse. Therefore, I suggest that in studies testing hypotheses related to the amount of intra-specific trait variation, the quadratic coefficient of variation should be used, especially if the inverse of the studied trait (i.e 1/trait) is also meaningful.

### 1. Materials and Methods

An R function for calculating two estimates of CV (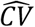 and 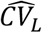), geometric coefficient of variation (GCV), and quartile coefficient of variation (*CV*_*Q*_) were developed (Appendix B). All analyses were done in R environment, and the script and data will be available in a public repository.

For illustrating purposes, the dataset of Gyalus et al. (in press) was used that contains plot level measurement of leaf traits. In this paper, only specific leaf (SLA, leaf area in cm^2^ per leaf dry mass in g) data were used. Leaf mass per area (LMA) was calculated as 1/SLA. Four indices of relative variation of SLA and LMA were calculated for each species with at least 10 SLA data. Then the absolute differences between SLA and LMA in relative within-species variation and species rank according to within-species variation were calculated. Since 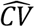 could be more sensitive to outlier values than other measures, all analyses were repeated after excluding outlier values.

## Acknowledgement

This research was supported by the NKFIH-K124671 grant.

## Data availability

Data and code available from Zenodo DOI 10.5281/zenodo.6907699.

## Authors’ contribution

ZB-D conceived, designed, and executed this study and wrote the manuscript. No other person is entitled to authorship

